# Spermidine suppresses DC activation via eIF5A hypusination and metabolic adaptation

**DOI:** 10.1101/2023.12.07.570665

**Authors:** Gavin R Meehan, Utku Gunes, Hannah E Scales, George Finney, Ross Deehan, Sofia Sintoris, Aegli Athanasiadou, James M Brewer

## Abstract

Cell metabolism plays an important role in immune effector responses and through responding to metabolic signals, immune cells can adapt and regulate their function. Arginine metabolism in Dendritic cells (DC) has been shown to reduce T cell activation; however, it is unclear how this immunosuppressive state is induced. To address this issue, we examined the immunomodulatory capacity of various metabolites from arginine metabolism. Through the use of a recently described DC:T cell interaction assay and flow cytometry we demonstrated that spermidine most significantly inhibited DC activation, preventing subsequent interactions with CD4 T cells. DC function could be restored by addition of inhibitors of spermidine metabolism via the eIF5A-hypusine axis, required for expression of some mitochondrial enzymes. We also demonstrated that the spermidine induced-immunosuppressive state protected DC against activation induced loss of mitochondrial capacity for energy generation, which was also hypusination dependent. Taken together this data demonstrates that spermidine is the key immunomodulatory component downstream of arginine metabolism and that it mediates this effect by stimulating hypusination-dependent protection of OXPHOS in DC, which in turn results in a reduced ability of DC to activate and interact with T cells. This pathway may be utilised by the immune system to regulate excessive immune responses but could also be exploited by pathogens as a method of immune evasion.

## Introduction

Cellular metabolic pathways such as Glycolysis, Oxidative Phosphorylation (OXPHOS), Fatty Acid Oxidation/Synthesis and amino acid metabolism are increasingly implicated in regulating activation and proliferation of immune cells^1–3^. While a wide range of metabolic changes are required to support the bioenergetic demands of cell activation, many metabolic systems are more closely involved in selectively promoting effector phenotypes underlying immunity and disease^4^. For example, Arginine metabolism has been widely studied in macrophage activation, where classically activated (M1 or inflammatory) macrophages upregulate nitric oxide synthase (NOS2) and metabolise arginine to produce nitric oxide as an antimicrobial agent^5^. In contrast, alternatively activated (or M2) macrophages express Arginase, leading to polyamine and collagen biosynthesis and tissue repair^5,6^.

Dendritic cells (DC) are professional antigen presenting cells with the unique ability to activate naïve T cells, as well as influence the magnitude and phenotype of the developing T cell response. They perform this role by sensing and responding to environmental stimuli such as pathogen- and danger-associated molecular patterns (PAMPs and DAMPs). These activating stimuli induce metabolic changes in DC that play an important role in supporting the effector role of DC in initiating T cell responses^7^. For example, TLR activation of DC stimulates an increase in glycolysis and mitochondrial oxidative phosphorylation to meet the energetic and biosynthetic demands of T cell activation^7^. Later in the response, inflammatory dendritic cells metabolise arginine via nitric oxide production which acts to block mitochondrial respiration requiring a shift toward glycolysis for ATP synthesis^7^. In contrast, under certain conditions, arginase driven metabolism of arginine results in reduced T cell activation, potentially contributing to immune tolerance or suppression of excessive immune responses^5^. The role of the arginase pathway in mediating reduced T cell activation is thought to be through the depletion of L-arginine, which is an essential amino acid required for T cell proliferation and function^5,8^. However, more recent studies demonstrate potentially immune modifying activity for downstream products of the arginase pathway, for example, polyamines in modifying mitochondrial respiration^9^ and *γ*-aminobutyric acid (GABA) in driving autophagy for intracellular remodelling^10^ required to support T cell activation.

Here we investigate the immunomodulatory activity of downstream metabolites of the arginase pathway. We demonstrate that exogenously added spermidine had 100 to 1000-fold greater inhibition of the ability of DC to interact with T cells compared with other products of arginase metabolism, which correlated with the ability of spermidine to inhibit DC activation. The inhibitory effects of spermidine on DC activation and T cell interaction were overcome by addition of GC7, demonstrating that the spermidine eIF5A-hypusine axis was responsible for immunomodulation. Together with subsequent metabolic analysis, our data demonstrate that exogenous spermidine stimulates spares activation induced loss of mitochondrial OXPHOS for energy generation, leading to suppression of DC effector functions and reduced T cell responses.

## Methods

### Animals

C57BL/6 female mice (6–10-week-old) were purchased from Envigo (Wyton, United Kingdom). Ovalbumin (OVA) peptide (323-339)-specific TCR transgenic (CD45.1^+^ OTII) mice on a C57BL/6 background (6–10-week-old) were produced in-house. All animals were maintained on a 12-hour light/dark cycle and provided with food and water ad libitum. All work was carried out under a UK Home Office licence in accordance with the Animals (Scientific Procedures) Act 1986 at the Central Research Facility at the University of Glasgow.

### Metabolites and Inhibitors

Stock solutions of L-arginine (Sigma-Aldrich), L-ornithine (Sigma-Aldrich), agmatine (Sigma-Aldrich), spermidine (Sigma-Aldrich), putrescine (Sigma-Aldrich), GABA (Tocris), DENSPM (Sigma-Aldrich), GC7 (Sigma-Aldrich), and CPX (Sigma-Aldrich) were freshly prepared and diluted to given concentrations in complete media which consisted of RPMI-1640 medium (ThermoFisher; Waltham, MA, USA) supplemented with 2 mM L-glutamine (Sigma-Aldrich; St Louis, MO, USA) ), 1x penicillin-streptomycin (Sigma-Aldrich; St Louis, MO, USA) and 10% foetal bovine serum (FBS) (ThermoFisher; Waltham, MA, USA). The pH of 50 mM L-arginine was adjusted to pH7 with concentrated hydrochloric acid prior to dilution .

### Dendritic Cell Culture

Bone-marrow derived dendritic cells (BMDCs) were produced as described previously^11^. Briefly, long bones were collected from C57BL/6 mice and flushed with complete media. Cells were seeded in 12 well plates (Corning) at a density of 7.5×10^5^ cells/ml in 2 ml complete media supplemented with 20 ng/ml recombinant human GM-CSF (Peprotech). The plates were incubated at 37°C in 5% CO_2_. On day 3, 1 ml of media supplemented with 20 ng/ml recombinant human GM-CSF (Peprotech) was added to each well. On day 6 half the media was removed and replaced from each well. The BMDCs were ready for use on day 7.

### Dendritic Cell Metabolite Cultures

On day 7, all the media was removed from the BMDCs and replaced with fresh complete media supplemented with spermidine with or without different metabolites and inhibitors. The specific concentrations of metabolites and inhibitors are indicated in the figure legends. The cells were incubated at 37°C in 5% CO_2_. 24 hours later 20ng/ml of LPS or media only was added to each well. The plates were incubated at 37°C at 5% CO_2_ for a further 24 hours and were subsequently processed for flow cytometry.

### CD4 T Cell Enrichment

Lymph nodes and spleens were collected from CD45.1 OTII mice and processed into single cell suspensions. The spleens were incubated with eBioscience RBC lysis buffer (Thermofisher; Waltham, MA, USA) for 5 minutes at room temperature and washed twice with complete media. The CD4+ T cells were then enriched by negative selection using a CD4 T cell isolation kit as per manufacturer’s instructions (Miltenyi Biotec; Bergisch Gladbach, Germany).

### Flow Cytometry

Plates of BMDCs were chilled on ice for 10 minutes before cells were harvested using a cell scraper. The cells were washed in Ca^2+^/Mg^2+^ free PBS containing 2mM EDTA at x400*g* for 5 minutes at 4°C and then resuspended in 400μl of Ca^2+^/Mg^2+^ free PBS containing eFluor506 or eFluor780 fixable viability dye (1:1000) (Thermofisher; Waltham, MA, USA) for 20 minutes. The cells were washed again in Ca^2+^/Mg^2+^ free PBS containing 2mM EDTA at x400*g* for 5 minutes at 4°C and resuspended in 100μl conditioned media from the anti-CD16/CD32 antibody producing hybridoma (2.4G2) (Fc Block). Antibody suspensions were prepared consisting of Fc Block and either MHCII-e450 (1:400), CD80-PE-Cy7 (1:100), CD86-FITC (1:100), CD40-PE (1:100) and CD11c-PerCP-Cy5.5 (1:100) or MHCII-APC-Cy7 (1:400), CD80-FITC (1:100), CD86-PE-Cy7 (1:100), CD40-APC (1:100) and CD11c-e450 (1:100) (all antibodies from Biolegend, San Diego, CA, USA). Cells were incubated with 100μl of the antibody suspension for 20 minutes at 4°C before a final wash in Ca^2+^/Mg^2+^ free PBS containing 2mM EDTA at x400*g* for 5 minutes at 4°C. The cells were resuspended in 200μl Ca^2+^/Mg^2+^ free PBS containing 2mM EDTA before analysis on a BD LSR fortessa flow cytometer (BD Biosciences, San Jose, USA). Analysis of flow cytometry data was performed using FlowJo 10.8.1 (FlowJo LLC, Ashland, OR, USA).

### DC - T Cell Interaction Assay

DC and T Cell co-cultures were setup and analysed using an INCell Analyzer 2000 (GE) as described previously^12^. Briefly, DCs and CD4+ T cells were washed and resuspended in RPMI supplemented with 2% FCS at a cell density of 1×10^7^ cells/ml and labelled with 7.5μM Cell Tracker Red CMTPX (Thermofisher; Waltham, MA, USA) or 7.5μM vibrant CFDA SE (CFSE) cell tracer kit (Thermofisher; Waltham, MA, USA) respectively. The cells were incubated at 37°C for 10 minutes and then washed 3x in complete media at x400g for 5 minutes at 4°C.

The cells were mixed 1:1 and resuspended in complete media. They were then seeded in 384-well μ-clear tissue culture treated microplates (Greiner; Kremsmünster, Austria) at a density of 16,000 cells/well. Cells were incubated in the presence of either 1 μg/mL OVA peptide (323-339; pOVA) (Sigma-Alrdrich; St Louis, MO, USA), 2.5 μg/mL concanavalin A (ConA) (Sigma-Alrdrich; St Louis, MO, USA) or with media only. As specified in the figure legends, various metabolites were also added to the wells. The plates were incubated at 37°C at 5% CO_2_ for 18 hours to assess changes in the percentage of DC: T cell overlap as a measure of cellular interaction.

Images were acquired using an INCell Analyser 2000 (GE Healthcare, Chicago, IL, USA) with the following settings: 10x magnification; 0.45 numerical aperture; flat field and apochromatic corrections (CD160) and chromatic aberration free infinity (planApo). Cells were imaged at 37°C with FITC captured using a 490/20 nm excitation and 525/36 nm emission filter while CMTPX was captured using 555/25 nm excitation and 605/52 nm emission filter. Each image captured 25% of the well and the percentage overlap was calculated using the Developer Toolbox software V1.9.2 (GE Healthcare, Chicago, IL, USA).

### Analysis of Cellular Metabolism

BMDCs were seeded in uncoated Seahorse XFp cell culture miniplates (Agilent, Santa Clara, CA, USA) at a density of 8 × 10^4^ cells/per well in complete media and incubated for 24 hours at 37°C in 5% CO_2_. Spermidine and GC7 were added to a final concentration of 0.1 mM and 1 μM respectively and the plates were incubated for a further 24 hours at 37°C in 5% CO2. LPS (100 ng/mL) or the equivalent volume of media was added to each well and the plates were incubated for a final 24 hours at 37°C in 5% CO2. The oxygen consumption rate (OCR) and the extracellular acidification rate (ECAR) were measured with an XF Cell Mito Stress Kit (Agilent, Santa Clara, CA, USA) as per manufacturer’s instructions. The kit used three modulators of mitochondrial phosphorylation which were oligomycin (1.5 mM), carbonyl cyanide 4-(trifluoromethoxy) phenylhydrazone (FCCP) (1 mM), and rotenone/antimycin A (1 mM). Prior to initiation of the assay the media was removed and replaced with XF RPMI Base Medium supplemented with 1 mM pyruvate, 2 mM glutamine and 10 mM glucose (All from Agilent, Santa Clara, CA, USA). The plates were read using a XF HS Mini analyser (Agilent, Santa Clara, CA, USA) and the data was analysed using the accompanying software.

### Statistical Analysis

All graphs and statistical analyses were produced using GraphPad Prism 9 (GraphPad Software Inc., San Diego, CA, USA). P values < 0.05 were deemed to be significant.

## Results

To identify which arginine metabolites had immunosuppressive effects, we performed a high throughput DC-T interaction assay which examined the effect that different metabolites had upon DC-T cell interactions. GABA (Figure 1A) was found not to influence ConA or pOVA driven DC-T cell interactions at any concentration. In contrast, ornithine (Figure 1B), putrescine (Figure 1C), arginine (Figure 1D) and agmatine (Figure 1E) significantly reduced ConA and pOVA induced DC-T cell interactions with metabolite concentrations between 5mM-50mM and 10mM-50mM (One Way ANOVA, P<0.05). Arginine and ornithine also significantly reduced interactions at lower concentrations with ConA, however this was not a consistent response. Spermidine (Figure 1F) had the greatest impact on DC-T cell interactions, producing a significant dose-dependent reduction at all concentrations (One Way ANOVA, P<0.05).

**Figure 1.**
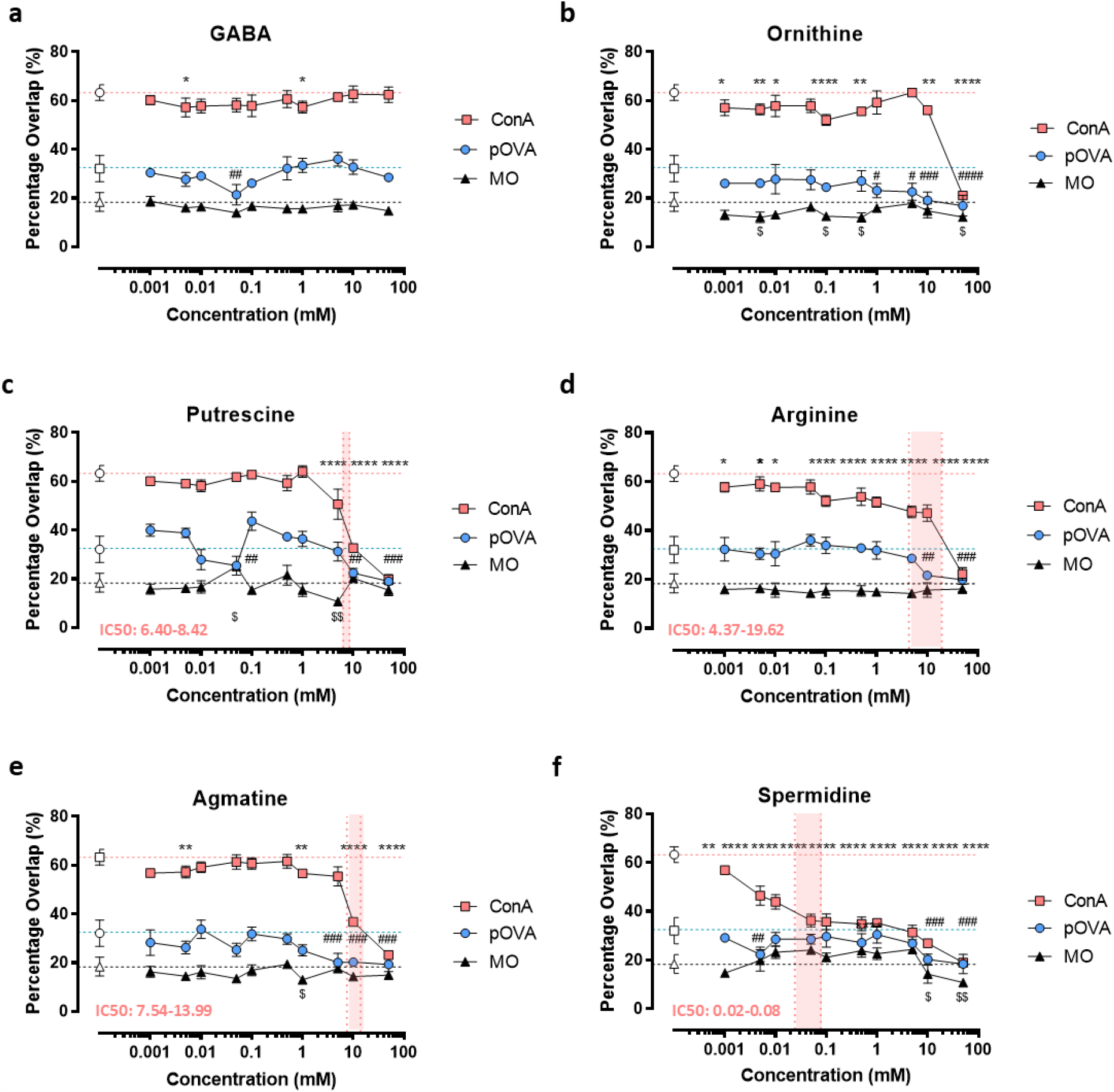
Exogenous arginine pathway metabolites alter DC-T Cell interactions *in vitro*. T cell and dendritic cell (DC) co-cultures were incubated overnight with various concentrations (50mM – 1μM) of different arginine pathway metabolites; (a) GABA, (b) ornithine, (c) putrescine, (d) arginine, (e) agmatine and (f) spermidine in the presence of either 1 μg/mL pOVA, 2.5 μg/mL ConA or media only (MO). Images were taken of each well using an INCell 2000 analyser and the percentage overlap between DCs and T cells was used to determine the level of cellular interaction. IC50 ranges, when able to be calculated, are shown for ConA in red. n = 3. One Way ANOVA; *,#, $, p < 0.05; **, ##, $$ p<0.01; ***, ### p< ***, p <0.001; ****, #### p<0.0001. * denotes statistical significance between ConA + metabolites (filled squares) vs ConA only (open squares). # denotes statistical significance between pOVA + metabolites (filled circles) vs pOVA only (open circles). $ denotes statistical significance between MO + metabolites (closed triangle) vs MO control (open triangle).

We next performed a series of flow cytometry experiments to determine if the effect of spermidine on DC-T cell interactions was associated with changes in DC activation. Based upon the IC50 calculated from the DC-T interaction assay, we incubated BMDCs with 0.1mM spermidine in the presence of the TLR4 agonist LPS. We demonstrated a significant reduction in the percentage of cells positive for the activation markers CD40 (Figure 2A) and CD86 (Figure 2B) (One Way ANOVA, P<0.05) while no effect was observed on the percentage of cells expressing CD80 (Figure 2C) and MHCII (Figure 2D). Assessment of the median fluorescent intensity (MFI) of these markers indicated a similar pattern with significantly reduced expression of CD40 (Figure 2E), CD80 (Figure 2B) and CD86 (Figure 2G) whilst MHCII was found to be unaffected (One Way ANOVA, P<0.05).

**Figure 2.**
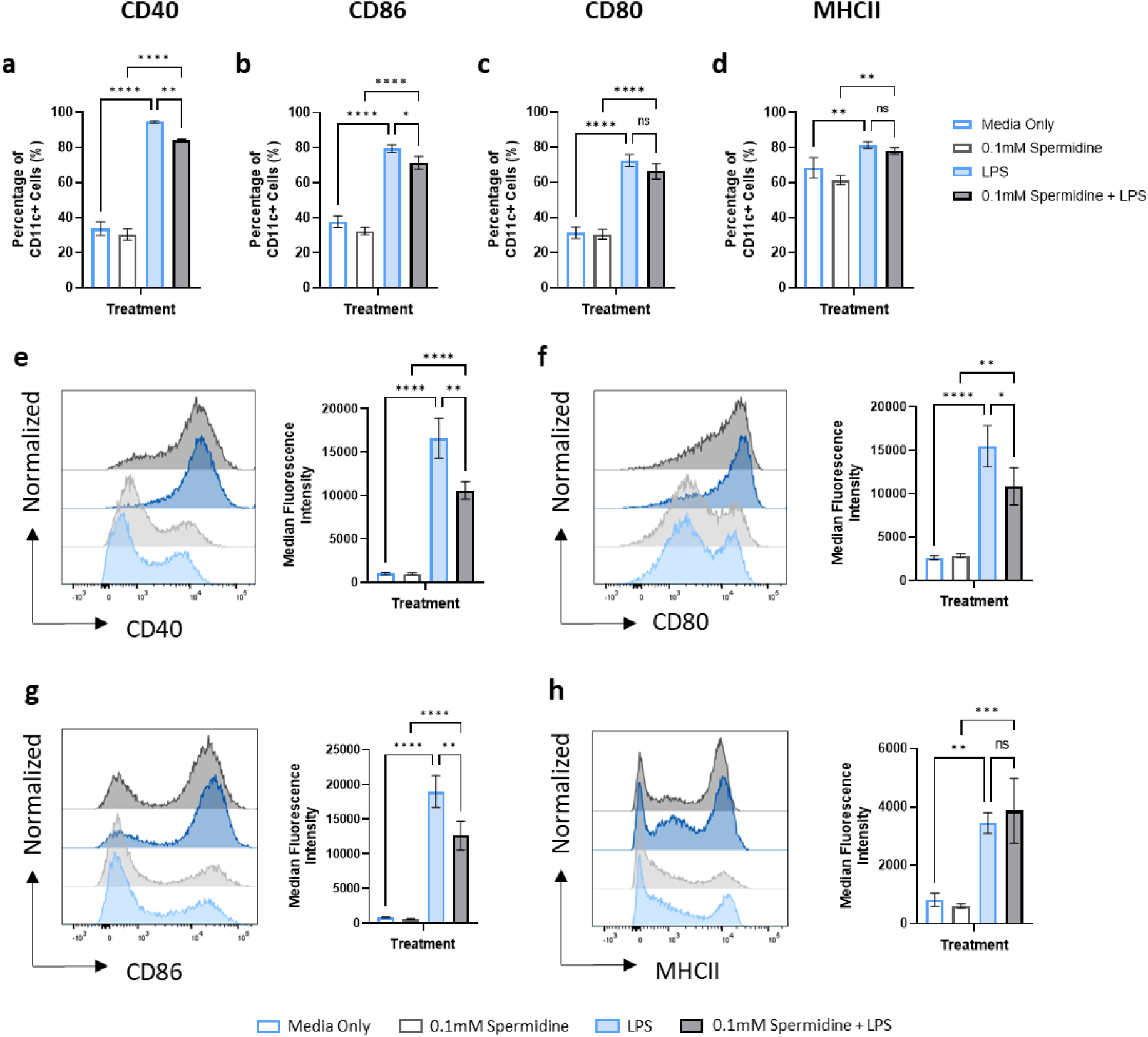
Spermidine Treatment inhibits BMDCs Function. Bone marrow derived dendritic cells (BMDCs) were incubated with or without 0.1mM spermidine in the presence of 20ng/ml LPS or media only for 24 hours. The BMDCs were then harvested, stained and analysed by flow cytometry. The percentage of cells expressing (a) CD40, (b) CD80, (c) CD86 and (d) MHCII were compared between the different treatment groups. The median fluorescent intensities (MFI) of (e) CD40, (f) CD80, (g) CD86 and (h) MHCII were also assessed between the treatment groups. n=3. One Way ANOVA, ns = no significance, *<0.05, **<0.01, ***<0.001, ****<0.0001

Spermidine is synthesised via the polyamine pathway where it acts a precursor for spermine synthesis and as a substrate for hypusination of eIF5A (Figure 3A). To determine how spermidine induces immunosuppression we utilised a number of analogues that interfere with the polyamine pathway. These included DENSPM, which promotes catabolism of spermine and spermidine; GC7 which inhibits synthesis of an eIF5A intermediate and CPX which inhibits eIF5A hypusination. Using the DC-T interaction assay, we incubated BMDCs with these inhibitors in the presence of ConA and 0.1mM spermidine. We found that DENSPM was unable to reverse spermidine induced inhibition of DC-T cell interactions at any concentration (Figure 3B). In contrast CPX was shown to have an effect at high concentrations (Figure 3C) whilst GC7 was found to have a significant dose-dependent effect at all concentrations (Figure 3D) (One Way ANOVA, P<0.05) producing an EC50 between 0.43 and 2.39μM.

**Figure 3.**
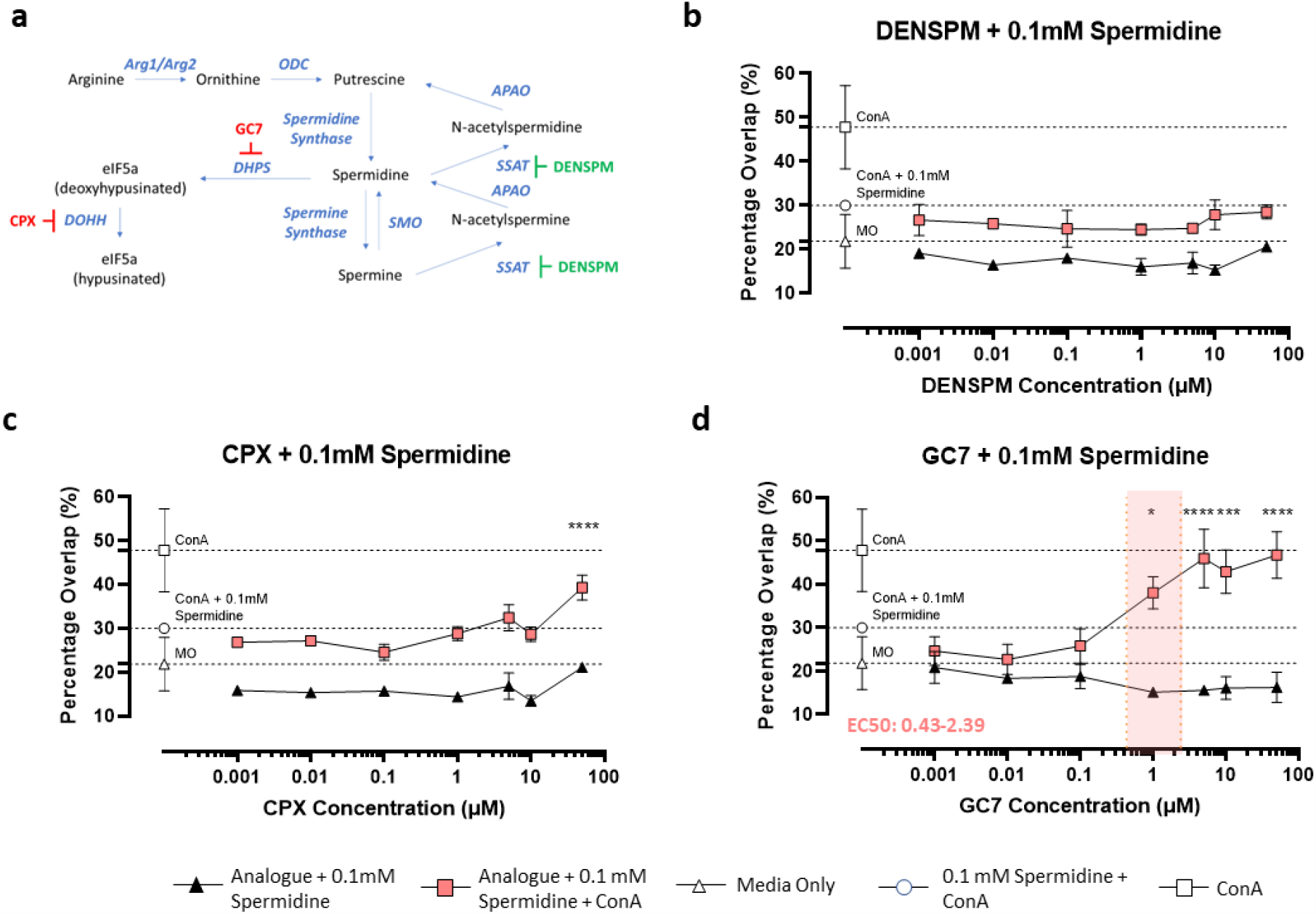
Polyamine analogues reverse spermidine induced inhibition of DC-T cell interactions *in vitro*. (a) Spermidine is synthesised via the polyamine pathway and acts as a precursor for spermine and as a substrate for the hypusination of eIF5a. The polyamine analogue DENSPM induces expression of SSAT and promotes polyamine catabolism and the polyamine analogues GC7 and CPX inhibit DHPS and DOHH respectively blocking eIF5a hypusination. (b-d) T cell and dendritic cell (DC) co-cultures were incubated overnight with various concentrations (50μM – 0.1nM) of different polyamine analogues: (b) DENSPM; (c) CPX; and (d) GC7 in the presence of 0.1mM spermidine with or without the addition of 2.5 μg/mL ConA. Images were taken of each well using an INCell 2000 analyser and the percentage overlap between DCs and T cells was used to determine the level of cellular interaction. n = 3. One Way ANOVA; * < 0.05; ***<0.001; ****<0.0001. * denotes statistical significance between ConA + 0.1mM spermidine + polyamine analogues (filled symbols) vs ConA + 0.1mM spermidine only (open symbols). EC50 ranges, where possible to calculate, are shown in red.

To confirm the results from the DC-T interaction assay and show definitively that the immunosuppressive effect of spermidine was mediated by eIF5A hypusination in DC, we performed further flow cytometry experiments where BMDCs were incubated with spermidine and 1μM GC7 in the presence of LPS. These experiments confirmed that GC7 treatment reversed spermidine induced inhibition with a significant increase in the proportion of cells expressing the activation markers CD40 (Figure 4A), CD80 (Figure 4B), CD86 (Figure 4C) and MHCII (Figure 4D) compared to spermidine treatment alone (One Way ANOVA, P<0.05). A similar increase was also found with the MFIs of these markers with CD40 (Figure 4E), CD80 (Figure 4F), CD86 (Figure 4G) and MHCII (Figure 4H) all being expressed in significantly higher levels in the GC7 treated cells (One Way ANOVA, P<0.05).

**Figure 4.**
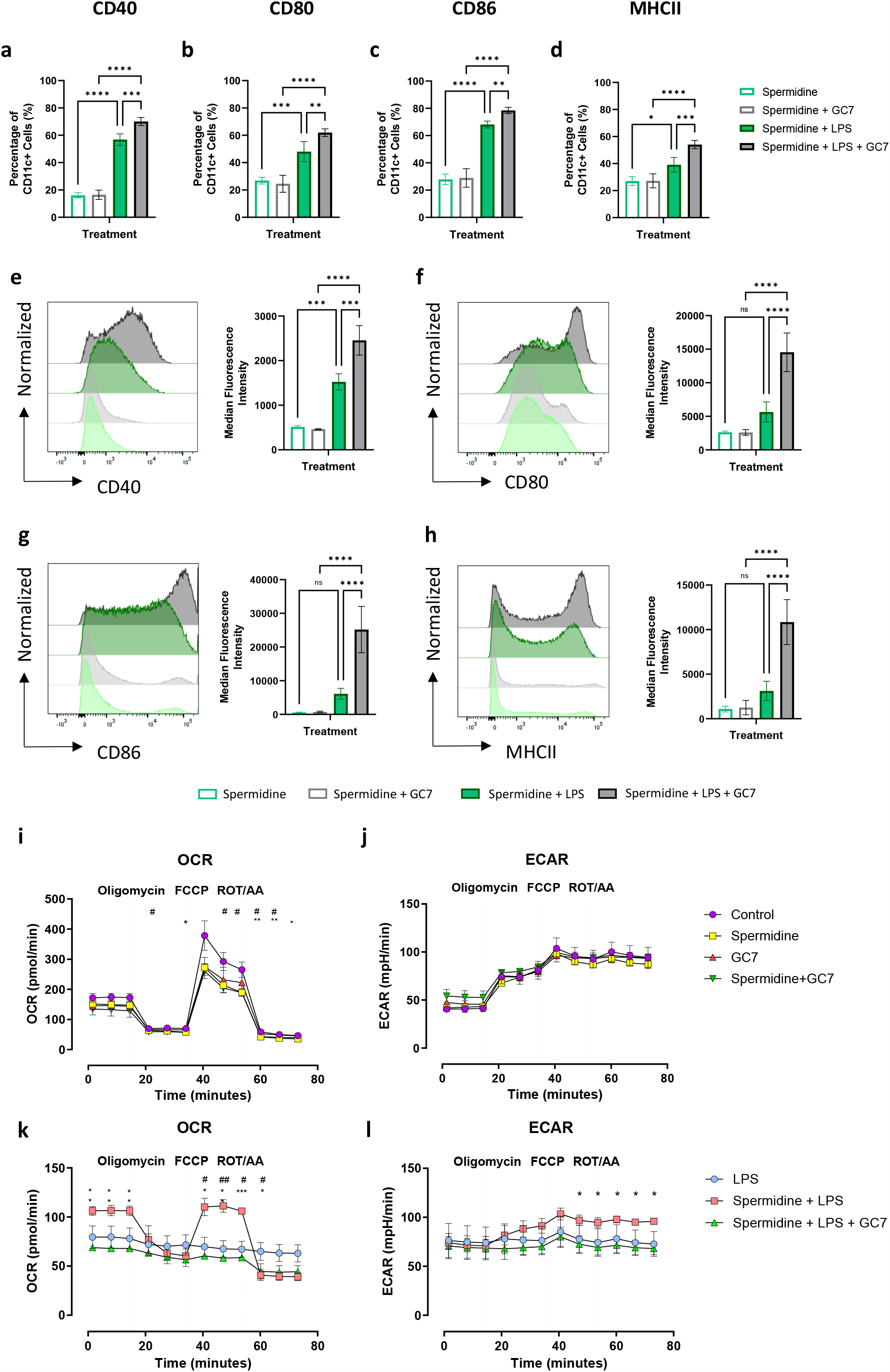
Spermidine analogue GC7 Prevents Spermidine Induced Inhibition of BMDC Function. Bone marrow derived dendritic cells (BMDCs) were incubated with 0.1mM spermidine with or without 1μM GC7 in the presence of 20ng/ml LPS or media only for 24 hours. The BMDCs were then harvested, stained and analysed by flow cytometry. The percentage of cells expressing (a) CD40, (b) CD80, (c) CD86 and (d) MHCII were compared between the different treatment groups. The median fluorescent intensities (MFI) of (e) CD40, (f) CD80, (g) CD86 and (h) MHCII were also assessed between the treatment groups. n=3. One Way ANOVA, ns = no significance, *<0.05, **<0.01, ***<0.001, ****<0.0001. Metabolic analysis of BMDCs treated with LPS, 0.1mM spermidine + LPS, 0.1mM spermidine + GC7 + LPS, media only, 0.1 mM spermidine, GC7 and 0.1 mM spermidine + GC7, was performed to compare the (i and j) oxygen consumption rate (OCR) and the (k and l) extracellular acidification rate (ECAR) of the cells. n=3. Two Way ANOVA, (i and j) *<0.05, **<0.01. * denotes statistical significance between media only vs 0.1mM spermidine. # denotes significance between media only vs 0.1mM spermidine + GC7; (k and l) *<0.05, **<0.01, ***<0.001. * denotes statistical significance between 0.1mM spermidine + LPS vs 0.1mM spermidine + LPS + GC7. # denotes significance between 0.1mM spermidine + LPS vs LPS.

To better understand the impact of spermidine on cellular respiration we also examined the mitochondrial oxygen flux (OCR) and extracellular acidification rate (ECAR) of the cells under various conditions. While spermidine and/or GC7 treatment had a minor effect on OCR (Figure 4I) there was no impact on ECAR (Figure 4J). However, on activation of BMDC with LPS, spermidine pretreatment prevented loss of both the OCR (Figure 4K) and the ECAR (Figure 4L) compared with LPS alone, with GC7 treatment reversing this effect. For OCR, this is most clearly seen at baseline respiration, prior to oligomycin treatment, and at maximal respiration rate seen following addition of the uncoupling agent FCCP. Taken together, this data demonstrates that the effects of spermidine on both DC function and metabolism are dependent on the hypusine/eIF5A axis as demonstrated by reversal on addition of GC7.

## Discussion

Induction of arginine metabolism via the arginase pathway has been associated with reduced DC activation^13^ however, the specific mechanisms responsible for mediating this process have remained unclear. Here, we demonstrate that spermidine is the key downstream product of this pathway, acting via the hypusine/eIF5A axis to spare DC OXPHOS on activation, leading to unresponsive DCs that fail to activate T cells, thus limiting the development of adaptive immune responses.

Alterations in arginine metabolism have been associated with immunosuppression in a number of different contexts including inflammation where low polyamine levels inversely correlate with higher inflammatory responses and cancer where high polyamine levels are associated with increased tumour growth and malignancy^14,15^. Spermidine has been implicated in inducing immunosuppressive states in monocytes, macrophages and DCs both in experimental models and in human disease^16–18^. However, using the deoxyhypusine inhibitor GC7 we have demonstrated definitively that spermidine exerts its effects in DCs by initiating eI5FA hypusination. Although hypusination has been shown to support alternative activation of macrophages^9^ to the best of our knowledge we are the first to define its role in supressing DC function.

Previous studies have demonstrated that hypusination supports alternative macrophage activation through production of certain mitochondrial proteins, leading to increased mitochondrial respiration^9^. While we did not observe an increase in OXPHOS on addition of spermidine, when BMDC were activated spermidine pretreatment demonstrated enhanced OXPHOS in DC, via a hypusination dependent process. Furthermore, blocking hypusination inhibits the metabolic effects of spermidine and concomitantly restores DC function. This suggests that spermidine blocks DC activation by altering DC mitochondrial capacity via hypusination of eI5FA. In line with the observations in our study, others have shown an association between OXPHOS and an anti-inflammatory state in immune cells including macrophages and DCs^19,20^.

Regulating or mimicking metabolites within specific immune cells is an ongoing area of research that offers a potential strategy for treating autoimmune diseases^21^ but is unclear in DCs whether immunosuppression is induced directly via hypusination or indirectly via increased mitochondrial respiration. Certain RNA viruses, including influenza, directly hypusinate eIF5a in stromal cells to prevent interferon production and thus promote viral replication^22^. This same mechanism may be induced by spermidine to prevent DC activation however it is also possible that hypusination alters mitochondrial capacity and the OXPHOS pathway which itself leads to an immunosuppressive state. Hypusinated eIF5A controls the switch to OXPHOS in macrophages and that inhibition limits the activity of the TCA cycle and the expression of many mitochondrial proteins^9,23^. This may also occur in DCs but to test this, future studies may wish to focus on examining the molecular mechanisms downstream of hypusination or on the induction of OXPHOS via unrelated pathways to determine whether immunosuppression can be induced by other means.

Our study focussed on the addition of spermidine exogenously; however, we hypothesise that specific products within the immune milieu would be able to stimulate the DCs to induce spermidine production intracellularly. It is also possible that other cells within the local microenvironment may produce spermidine to alter the activation state of bystander DCs as had been shown with the gut microbiota, which produce spermidine to prevent T cell activation^16^. As spermidine is found in significant quantities in the diet, it is also possible that consumption of the metabolite may influence the levels of inflammation in specific tissues as has been observed in a mouse model of colitis^24^.

Our work focussed on examining the effect of spermidine in an *in vitro* setting, but other studies have demonstrated the anti-inflammatory effects of spermidine *in vivo* including research with a mouse model of Alzheimer’s disease which demonstrated that low amounts of the compound were able to reduce neuroinflammation^25^. It may therefore be useful for future research to focus on examining the effect of spermidine in different inflammatory conditions to establish whether it may offer potential insights into new anti-inflammatory drugs. Similarly, compounds such as GC7, which can inhibit eIF5A hypusination may be useful in enhancing immune responses and overcoming the immunosuppressive states that are induced in cancers and in certain infections.

In summary, we show that spermidine is the key immunomodulatory component downstream of arginine metabolism and that it mediates this effect by stimulating hypusination-dependent alterations in OXPHOS in DC, which in turn results in a reduced ability of DC to activate and interact with T cells.

## Statements

### Arrive Guidelines

All animal research adhered to ARRIVE guidelines (https://arriveguidelines.org/arrive-guidelines).

### Data Availability

Data is available to readers upon request to the corresponding author.

## Acknowledgements

The authors gratefully acknowledge the Flow Core Facility for their support & assistance in this work. The work of U. Gunes was supported by the Republic of Turkey Ministry of National Education(MoNE) under the scholarship MoNE-1416/YLSY.

## Author Contributions

GRM designed the research, constructed the figures, analysed the data and wrote the manuscript. UG performed flow cytometry experiments, metabolic analysis and analysed the data. HES designed the research, performed data analysis and constructed figures. GF performed DC: T cell interaction assays. RD, SS and AA performed and analysed flow cytometry experiments. JMB designed the research and contributed to writing the paper. All authors edited the paper.

## Permission to Reproduce

This work may be reproduced with permission of the corresponding author.

## Clinical trial registration

N/A

## References

1. O’Neill, L. A. J. & Pearce, E. J. Immunometabolism governs dendritic cell and macrophage function. J. Exp. Med. 213, 15–23 (2016).

2. Pearce, E. J. E. L. & Pearce, E. J. E. L. Immunometabolism in 2017: Driving immunity: all roads lead to metabolism. Nat. Rev. Immunol. (2017) doi:10.1038/nri.2017.139.

3. Corrado, M. & Pearce, E. L. Targeting memory T cell metabolism to improve immunity. J. Clin. Invest. 132, (2022).

4. Mills, E. L. et al. Itaconate is an anti-inflammatory metabolite that activates Nrf2 via alkylation of KEAP1. Nature 556, 113–117 (2018).

5. Bronte, V. & Zanovello, P. Regulation of immune responses by L-arginine metabolism. Nat Rev Immunol 5, 641–54 (2005).

6. Minutti, C. M., Knipper, J. A., Allen, J. E. & Zaiss, D. M. W. Tissue-specific contribution of macrophages to wound healing. Semin. Cell Dev. Biol. 61, 3–11 (2017).

7. Everts, B. & Pearce, E. J. Metabolic control of dendritic cell activation and function: Recent advances and clinical implications. Front. Immunol. 5, 1–7 (2014).

8. Geiger, R. et al. L-Arginine Modulates T Cell Metabolism and Enhances Survival and Anti-tumor Activity. Cell 167, 829–842.e13 (2016).

9. Puleston, D. J. et al. Polyamines and eIF5A Hypusination Modulate Mitochondrial Respiration and Macrophage Activation. Cell Metab. 1–12 (2019) doi:10.1016/j.cmet.2019.05.003.

10. Kim, J. K. et al. GABAergic signaling linked to autophagy enhances host protection against intracellular bacterial infections. Nat. Commun. 9, (2018).

11. Inaba, K. et al. Generation of large numbers of dendritic cells from mouse bone marrow cultures supplemented with granulocyte/macrophage colony-stimulating factor. J. Exp. Med. 176, 1693–1702 (1992).

12. Scales, H. E. et al. A discovery pipeline for identification and in vivo validation of drugs that alter T cell/ dendritic cell interaction. bioRxiv 2022.04.11.487526 (2022) doi:10.1101/2022.04.11.487526.

13. Simioni, P. U., Fernandes, L. G. & Tamashiro, W. M. Downregulation of L-arginine metabolism in dendritic cells induces tolerance to exogenous antigen. Int. J. Immunopathol. Pharmacol. 30, 44–57 (2017).

14. Chia, T., Zolp, A. & Miska, J. Polyamine Immunometabolism: Central Regulators of Inflammation, Cancer and Autoimmunity. Cells vol. 11 (2022).

15. Lian, J. et al. The role of polyamine metabolism in remodeling immune responses and blocking therapy within the tumor immune microenvironment . Frontiers in Immunology vol. 13 (2022).

16. Carriche, G. M. et al. Regulating T-cell differentiation through the polyamine spermidine. J. Allergy Clin. Immunol. 147, 335–348.e11 (2021).

17. Yang, Q. et al. Spermidine alleviates experimental autoimmune encephalomyelitis through inducing inhibitory macrophages. Cell Death Differ. 23, 1850–1861 (2016).

18. Wawrzyniak, M. et al. Spermidine and spermine exert protective effects within the lung. Pharmacol. Res. Perspect. 9, e00837 (2021).

19. Wculek, S. K. et al. Oxidative phosphorylation selectively orchestrates tissue macrophage homeostasis. Immunity 56, 516–530.e9 (2023).

20. Kelly, B. & O’Neill, L. A. J. Metabolic reprogramming in macrophages and dendritic cells in innate immunity. Cell Res. 25, 771–784 (2015).

21. Pålsson-McDermott, E. M. & O’Neill, L. A. J. Targeting immunometabolism as an anti-inflammatory strategy. Cell Res. 30, 300–314 (2020).

22. Seoane, R. et al. eIF5A is activated by virus infection or dsRNA and facilitates virus replication through modulation of interferon production . Frontiers in Cellular and Infection Microbiology vol. 12 (2022).

23. Barba-Aliaga, M. & Alepuz, P. Role of eIF5A in Mitochondrial Function. International Journal of Molecular Sciences vol. 23 (2022).

24. Niechcial, A. et al. Spermidine ameliorates colitis via induction of anti-inflammatory macrophages and prevention of intestinal dysbiosis. J. Crohn’s Colitis jjad058 (2023) doi:10.1093/ecco-jcc/jjad058.

25. Freitag, K. et al. Spermidine reduces neuroinflammation and soluble amyloid beta in an Alzheimer’s disease mouse model. J. Neuroinflammation 19, 172 (2022).

